# Interfering with plant developmental timing promotes susceptibility to insect vectors of a bacterial parasite

**DOI:** 10.1101/2022.03.30.486463

**Authors:** Weijie Huang, Saskia A. Hogenhout

## Abstract

Multi-trophic parasites often modulate host behaviors and development to attract other hosts. However, whether these modulations are controlled by specific parasite genes and are adaptive remains debated. Insect-transmitted phytoplasma parasites take control of their plant host by turning them into ‘Zombie plants’, which feature spectacular vegetative organ proliferations and juvenilization. The phytoplasma effector SAP05 induces these symptoms by mediating the degradation of multiple plant SPL and GATA developmental transcription factors via recruiting the ubiquitin receptor RPN10. Here, we investigated if these SAP05-induced modulations impact the leafhopper vectors on which phytoplasmas depend for spread. SAP05 promotes leafhopper reproduction on *Arabidopsis thaliana* in a RPN10-dependent manner. Moreover, SAP05 homologs that degrade SPLs and both SPLs and GATAs promote leafhopper reproduction, whereas those that degrade only GATAs do not. Leafhoppers also produced more progeny on plants with misregulated *MiR156*, which suppresses SPL expression. SPLs control several developmental processes. Surprisingly, leaf color and shape changes and increased leaf numbers did not correlate to leafhopper progeny increases, whereas disturbing the plant phase transition processes did. Therefore, only specific phenotypes induced by a single parasite gene extend beyond one host to control another and promote host-host attractions of a multi-trophic parasite.

## Introduction

Molecular mechanisms underpinning parasite-enforced host modifications have remained largely uncharacterized, leading to unclarities of whether these phenotypes are adaptive (Herbison *et al*., 2018; Hughes & Libersat, 2019; Doherty, 2020). Phytoplasmas are obligate bacteria that mostly rely on both host plants and insect vectors for survival and transmission (Fig. 1a) (Hogenhout *et al*., 2008). For example, the Aster Yellows phytoplasma strain Witches’ Broom (AY-WB) colonizes multiple plant species, including *Arabidopsis thaliana*, and is transmitted by the polyphagous leafhopper *Macrosteles quadrilineatus*. AY-WB secretes a plethora of effector proteins to modulate plant processes (Sugio *et al*., 2011b). Effector SAP11 binds and destabilize plant TCP transcription factors, enhancing insect vector reproduction by suppressing plant defense hormone biosynthesis while promoting leaf crinkling and branching (Sugio *et al*., 2011a). Effector SAP54 binds and degrades plant MADS-box transcriptions factors, turning flowers into leaf-like structures (phyllody) (MacLean *et al*., 2011; MacLean *et al*., 2014). SAP54 also promotes insect colonization on host plants via unknown mechanisms that are independent of phyllody formation (Orlovskis & Hogenhout, 2016). Effector SAP05 concurrently destabilizes most plant SPL and GATA developmental regulators leading to the uncoupling of plant developmental phase transitions that, in turn, induces bushy plants with “witches’ broom” features and delays plant death upon infection with the phloem-colonizing phytoplasmas (Huang *et al*., 2021). However, it is unknown how SAP05 impacts plant susceptibility to the phytoplasma insect vectors. In all cases, it is not fully understood how the effector-induced plant morphological changes are related to the success of this insect-vectored pathogen. The phytoplasma disease pathosystem represents an excellent opportunity to study the genetic basis as well as the adaptiveness of extended phenotypes.

**Fig. 1.**
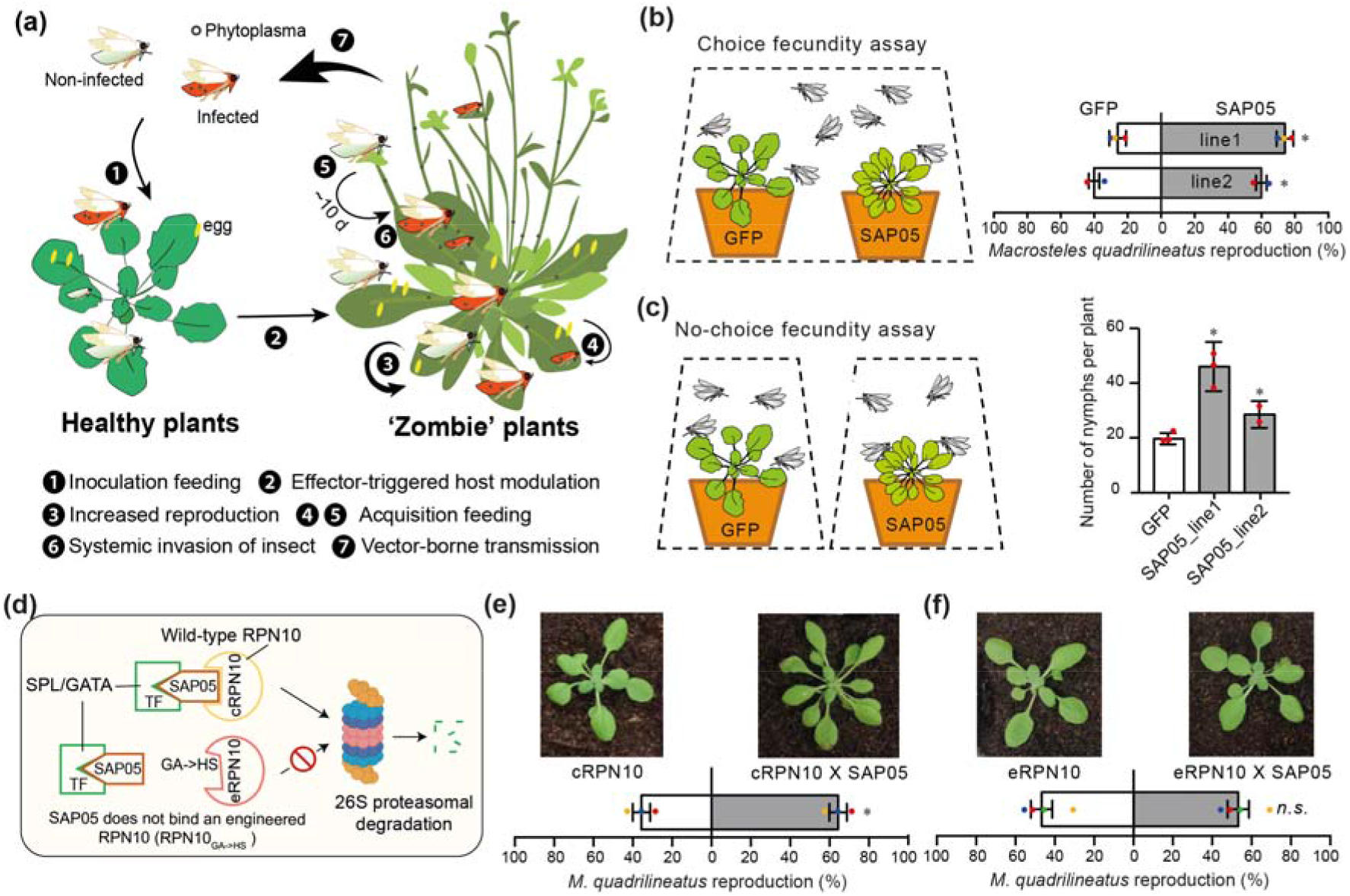
SAP05 promotes insect vector reproduction on *Arabidopsis*. (a) Lifecycle of the multi-trophic phytoplasma parasite AY-WB. The phytoplasmas are obligate plant pathogens that invasively colonize the major phloem-feeding insect vectors, *Macrosteles quadrilineatus* leafhoppers, within ± 10 days and depend on these insects for transmission to plants. Upon transmission to ± 4-week-old *Arabidopsis thaliana* plants, symptoms become apparent within about 14 days. The plants often exhibit increased leaf, stem proliferations and sterility and hence are dubbed ‘Zombie plants’. (b) SAP05 promotes *M. quadrilineatus* reproduction on *A. thaliana* in choice fecundity assay. Data are mean ± SD. \**P* < 0.05, χ^2^ test. (c) SAP05 promotes *M. quadrilineatus* reproduction on *A. thaliana* in no-choice fecundity assay. Data are mean ± SD. \**P* < 0.05, One-way ANOVA. (d) Diagram of how an engineered RPN10 allele confers *A. thaliana* resistance to SAP05 activities. (e, f) SAP05 promotes *M. quadrilineatus* reproduction on cRPN10 plants and not on eRPN10 plants. Plant photos show 4 weeks-old *A. thaliana* plants used in the assays. Data are mean ± SD. * *P* < 0.05, χ^2^ test.

## Materials and Methods

### Plant material and growth conditions

All *A. thaliana* genotypes used in the study were of Columbia-0 (*Col-0*) ecotype background. Plants were grown under 10 h light/14 h dark condition at 22°C on insecticide-free F2 compost soil (Levington).

### Rearing of insects

Phytoplasma-free aster leafhopper *M. quadrilineatus* colonies were reared on oat (*Avena sativa*) under 16 h light/8 h dark condition at 22°C.

### No-choice fecundity assay

For no-choice assays, individual *A. thaliana* of 4-5 weeks old was exposed to 6 female and 4 male adult *M. quadrilineatus* in a perforated bag. After 5 days, the insects were removed. The number of nymphs produced on each plant was scored 2-3 weeks after leafhopper removal. For each genotype, at least three plants were used in no-choice fecundity assays.

### Choice fecundity assay

For choice assay, three control plants and three experiment plants, placed in an alternating manner on a tray, were bagged together in a perforated bag. Twenty *M. quadrilineatus* (10 females and 10 males) were released into the bag for 5 days and removed. The plants were then bagged individually and the nymphs that developed on each plant were scored 2-3 weeks after leafhopper removal. The experiments were repeated two or three times.

### Phylogenetic analysis

Phylogenetic analysis was performed as in (Huang *et al*., 2021).

### Statistical Analysis

Statistical analysis was performed with Prism 7. χ^2^ test was used to analyze choice fecundity assay data. One-way ANOVA was used to analyze no-choice fecundity assay data. Two-tailed Student’s *t*-test was used for other analysis.

## Results

To investigate the impact of the SAP05-mediated plant host modulations on the performance of *M. quadrilineatus* leafhoppers, choice and no-choice fecundity experiments with transgenic *Arabidopsis* lines stably expressing SAP05 and GFP (control) were conducted. *M. quadrilineatus* produced more nymphs on SAP05 plants in both conditions (Fig. 1b,c). In a previous study we found that SAP05 does not bind an *A. thaliana* RPN10_GA>HS_ allele and, moreover, that the *rpn10 null* mutant complemented with this functional RPN10_GA>HS_ allele (eRPN10) does not develop witches’ broom-like symptoms in the presence of SAP05, as opposed to the *rpn10* mutants complemented with the wild-type RPN10 (cRPN10) (Fig. 1d) (Huang *et al*., 2021). Choice assays conducted herein showed that *M. quadrilineatus* produced more nymphs on wild-type RPN10 complementation (cRPN10) plants expressing *SAP05* (cRPN10 x SAP05) than on cRPN10 plants (Fig. 1e). In contrast, the leafhoppers produced similar numbers of nymphs on eRPN10 plants expressing *SAP05* (eRPN10 x SAP05) and eRPN10 plants (Fig. 1f). These data indicate that RPN10 is a susceptibility factor that SAP05 must interact with to promote the reproduction of insect vectors on plant hosts. Therefore, the SAP05-RPN10 interaction shapes not only the phytoplasma-plant interface, but also the next trophic level of the plant host interactions with phytoplasma insect vectors.

To investigate if SAP05-induced morphological changes are responsible for the increase in insect vector reproduction, we made use of both the *spl* and *gata* loss-of-function *A. thaliana* mutants that specifically phenocopy different features exhibited by the SAP05 transgenic *A. thaliana (*Huang *et al*., 2021). Because of redundancy, many single mutants from these gene families do not show obvious phenotypes in contrast to overexpressors (OEs) that exhibit clear morphological alterations. Therefore, several well-characterized overexpressors were also tested in choice fecundity assays to delineate the association between plant morphology and plant susceptibility to insects (fig. 2; fig. S1). Specifically, we performed the fecundity assays on 4-5 weeks old plants grown under short-day conditions, at a stage where most of the genotypes have not yet entered the reproductive stage and show distinct leaf developmental phenotypes. The plants remained vegetative throughout the assays.

**Fig. 2.**
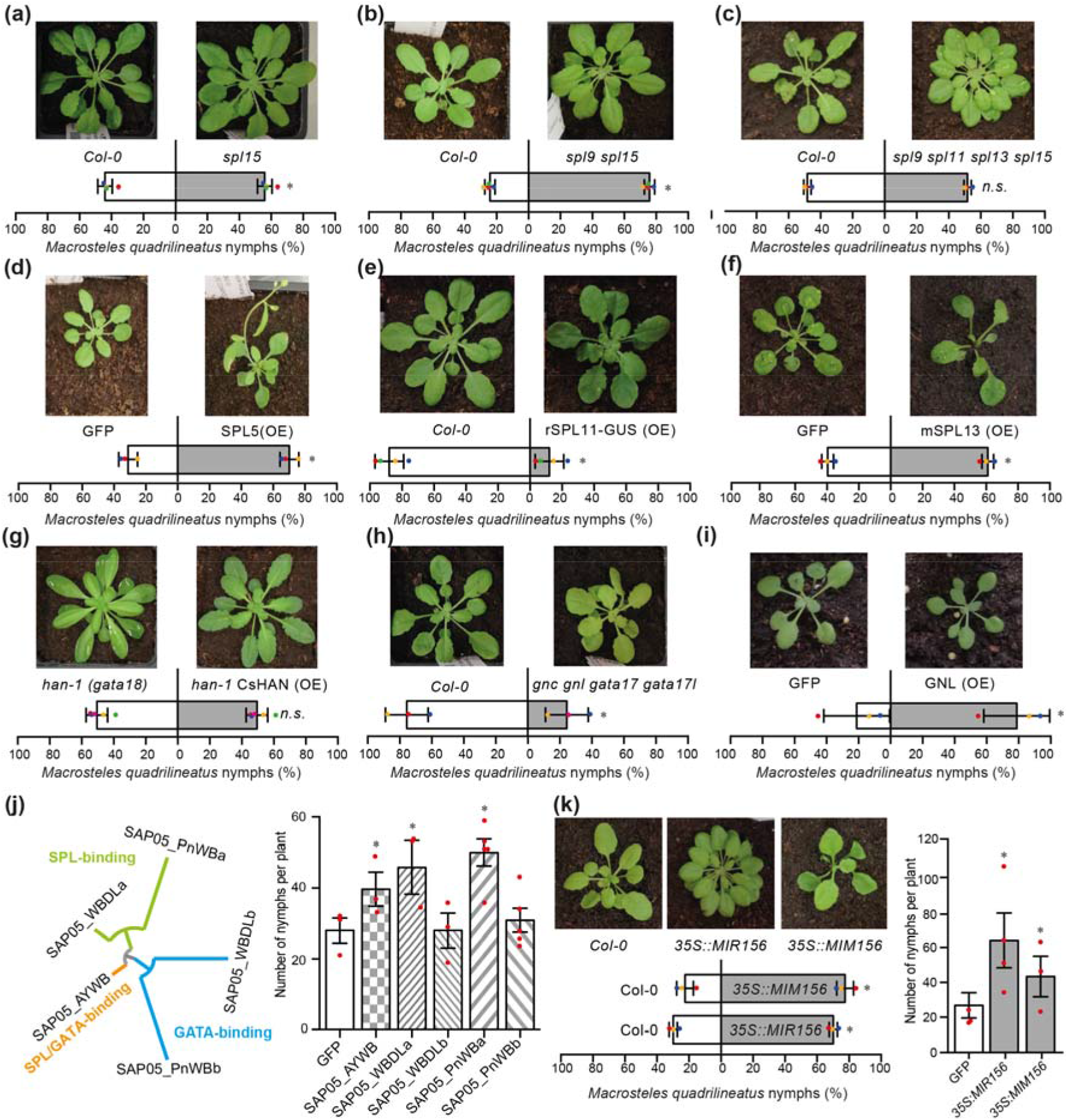
Specific SAP05-induced morphological changes promote phytoplasma insect vector performances on *A. thaliana*. (a-i) Choice fecundity assays on plants with misregulated levels of SPL or GATA family genes. Data are mean ± SD. * *P* < 0.05, χ^2^ test. (j) SPL-targeting SAP05 homologs promote *M. quadrilineatus* nymph production, whereas GATA-targeting SAP05 homologs did not. Data are mean ± SD. * *P* < 0.05, One-way ANOVA. (k) *M. quadrilineatus* nymph production is increased on plants with misregulated *MIR156*, which regulate transcript levels of SPL TFs. Data are mean ± SD. * *P* < 0.05, χ^2^ test in choice assays; One-way ANOVA in no-choice assays.

Given that SAP05 degrades most SPLs and GATAs, we hypothesized that *spl* and *gata* mutants are more susceptible to insects while *SPL* and *GATA* OEs are more resistant to insects in general. However, our results did not support this hypothesis. Even though *M. quadrilineatus* produced more nymphs on some *spl* mutants, including the *spl15* single mutant, the *spl9 spl15* double mutant and the *spl1 spl12* double mutant, these insects did not show a reproduction preference for the *spl9 spl11 spl13 spl15* quadruple mutant (Fig. 2a,b,c; Fig. S1a). Moreover, while the overexpression of *SPL11* negatively impacted *M. quadrilineatus* fecundity, the insects produced more nymphs on plants that overexpress *SPL5* or *SPL13* (Fig. 2d,e,f). On the GATA side, two *GATA* OEs, *GNC* and *GNL*, promoted insect nymph production choice while both the *gnc gnl* double mutant and the *gnc gnl gata17 gata17l* quadruple mutant had a negative impact (Fig. 2h,i; Fig. S1b,c). *GATA17* overexpression alone did not change insect nymph production (Fig. S1d). Therefore, SPL family and the GATA family members individually or in combination differentially shape the plant-insect interaction interface.

Visual cues, such as yellowing, have been shown to promote leafhopper attraction (Todd *et al*., 1990; Al-Subhi *et al*., 2021), it is possible that morphological attributes of the *spl* and *gata* mutants impacted leafhopper reproduction choice. The *spl9 spl11 spl13 spl15* quadruple mutant, the *gnc gnl* double mutant and the *gnc gnl gata17 gata17l* quadruple mutant all have yellower foliage (Fig. S1e). However, *M. quadrilineatus* either did not produce more progeny or even produced less progeny on those mutants (Fig. 2c,h; Fig. S1b). Therefore, in the *spl* and *gata* mutants tested, there appears to be no correlation between the level of yellowing and *M. quadrilineatus* reproduction success. Similarly, plant leaf numbers and sizes were also unlikely to have affected leafhopper reproduction preference. For instance, *SPL13* OE and *GNL* OE had significantly less leaves or smaller leaves (Fig. S1f,g) and were nonetheless preferred by *M. quadrilineatus* for nymph production (Fig. 2f,i). As well, the *spl9 spl11 spl13 spl15* quadruple mutant produced more leaves than the spl*15* or *spl9 spl15* mutants (Fig. S1h), but the insects did not produce more progeny on the quadruple mutant relatively to the single or double mutants (Fig. 2a,b,c). Furthermore, SAP05 plants had slender leaves with smooth margins, phenocopying *han-1* (*gata18*) plants while the overexpression of a *Cucumis sativus HAN* homologue in *han-1* background resulted in increased serration (Ding *et al*., 2015). However, *M. quadrilineatus* equally selected *han-2* and the *CsHAN* OE for nymph production (Fig. 2g). Therefore, the SAP05-induced increases in yellowing and leaf numbers and changes in leaf shapes do not seem to directly contribute to increased insect vector attraction and reproduction.

To differentiate the roles of SPL and GATA families in SAP05-mediated plant defense regulation, we took advantage of transgenic *A. thaliana* lines expressing SAP05 homologs that have evolved targeting specificities towards either SPLs or GATAs (Huang *et al*., 2021). Specifically, witches’ broom disease of lime (WBDL) phytoplasma and peanut witches’ broom (PnWB) phytoplasma, both have two copies of SAP05, with one copy (SAP05_WBDLa and SAP05_ PnWBa) targeting only SPLs and the other copy (SAP05_WBDLb and SAP05_PnWBb) targeting only GATAs (Fig. 2j). Those plants exhibited distinct leaf morphologies that resemble *spl* mutants and *gata* mutants, respectively, consistent with the targeting specificities of different SAP05 homologs (Huang *et al*., 2021). *M. quadrilineatus* produced more progeny in no-choice assays on SAP05_WBDLa and SAP05_PnWBa plants, whereas the progeny productions on SAP05_WBDLb and SAP05_PnWB plants were similar to that of wild type plants (Fig. 2j). In agreement with earlier data, *M. quadrilineatus* produced more progenies on SAP05_AYWB plants. Therefore, as opposed to specific single or higher *gata* mutants, the SAP05-targeting of multiple GATAs have a neutralizing effect on leafhopper fecundity choice, and the promotion of leafhopper reproduction choice can be largely explained by the SAP05-mediated targeting of multiple SPL members.

To further understand the roles of SPL in plant-insect interaction, we made use of the well-characterized *A. thaliana* lines that overexpress *MiR156*, which represses the expression of 10 *SPL* genes, or lines that express *MiM156*, which blocks MiR156 functioning (Wang *et al*., 2009). We found that the insects reproduced more progeny on both genotypes, regardless of choice and no-choice settings (Fig. 2k). Together with our previous finding that certain *spl* mutants and *SPL* OEs were both able to better support insect vector reproduction, our results indicate that proper developmental timing controlled by *MiR156* and downstream *SPL* genes, rather than the degradation of individual SPLs, enables *Arabidopsis* plants to mount appropriate defense responses to the phytoplasma insect vectors. Nevertheless, the degradation of SPLs by phytoplasma SAP05 likely reduce plant defense against the insect vector.

## Discussion

Whereas the miR156-SPL network has been primarily studied in the plant developmental context, evidence its roles in regulating plant-biotic interactions, including to insect herbivores, has been accumulating (Mao *et al*., 2017; Ge *et al*., 2018). Interestingly, it was shown that plant insect resistance exhibits age-regulated dynamics, with jasmonate response highly active at juvenile stage but declining with age while the accumulation of defense metabolite contributing to the insect resistance in adult plants (Mao *et al*., 2017). Our work here indicates that the interplay of multiple SPLs, rather than just one or a few, plays a role in promoting plant susceptibility to phytoplasma leafhopper vectors. We propose that trade-offs between growth and defense are integral components in the plant developmental transitions. As such, defects in the plant developmental phase transitions, leading to either juvenilization or premature aging, likely undermines plant defense mechanisms.

The effects of genes extending beyond individual organisms into environments are coined extended phenotypes (Dawkins, 1982). However, whether these parasite-enforced modifications are adaptive has been much debated. Our results herein showcase that a single gene of vector-borne parasite has evolved to modulate host developmental timing as a key virulence strategy, inciting both morphological and physiological changes in plants that collectively confer adaptive advantages for this ‘Zombie plant’ pathogen. Among those changes, some promote pathogen colonization, some enhance the vector performance, and some may not benefit the pathogen. Therefore, extended phenotypes can be complex, and a better understanding of their genetic basis holds the key to assess the adaptiveness of host modulations.

## Acknowledgements

We thank the JIC Entomology Facility for maintaining leafhopper and phytoplasma stocks; the John Innes Centre (JIC) Horticultural Services for growing plants; and JIC Photographic Services for imaging plant materials. This work was funded by the Human Frontier Science Program RGP0024/2015, Biotechnology and Biological Sciences Research Council grants BBS/E/J/000PR9797 and the John Innes Foundation.

## Author Contribution

S.A.H. obtained funding. W.H. and S.A.H. designed the approach. W.H. conducted the experiments. W.H. and S.A.H. wrote the manuscript.

## Data availability

All data are included in the article and SI Appendix.

---

**Fig. S1.**
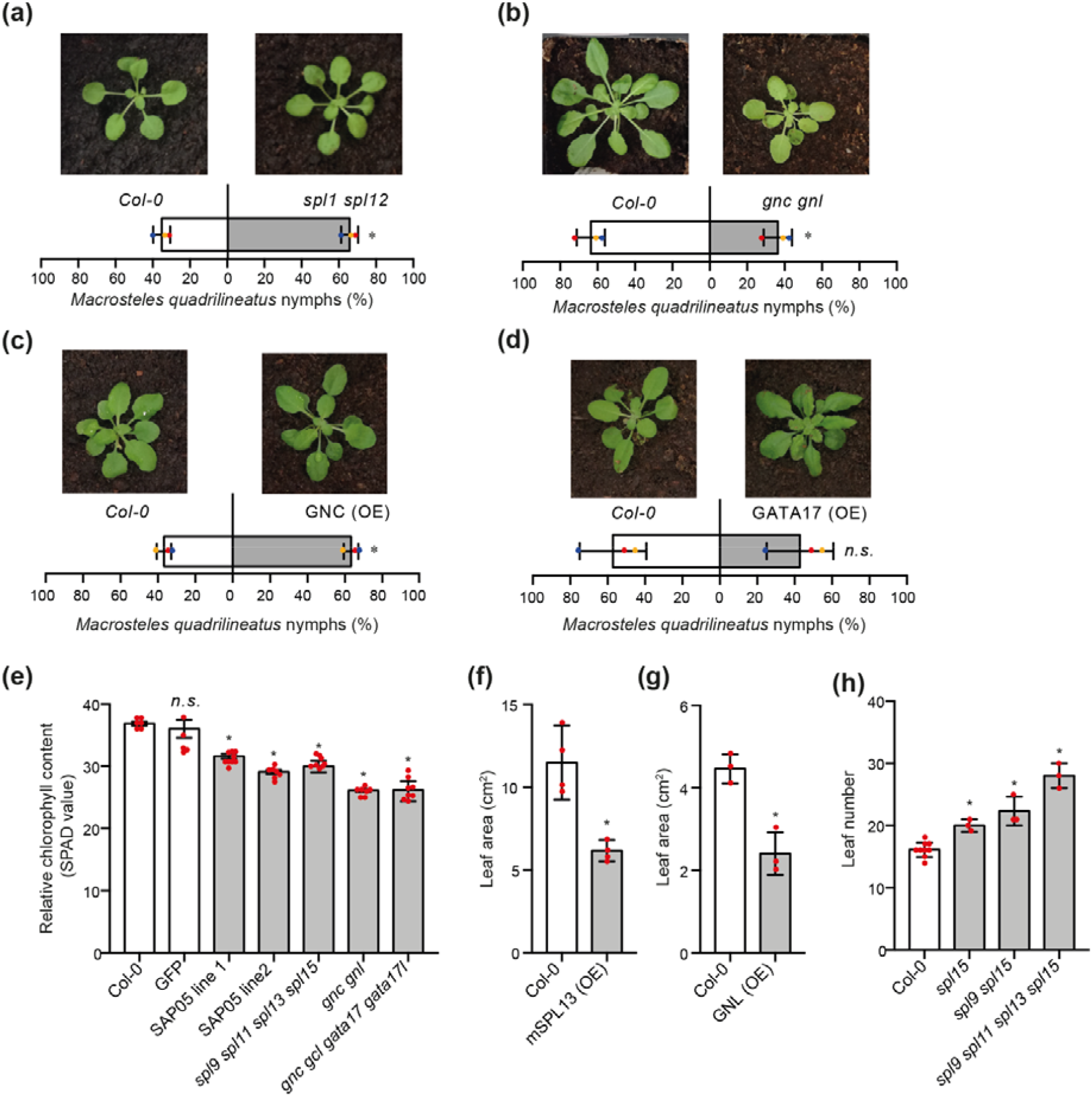
Choice fecundity assay results and the quantification of leaf development on plants used in the study. (a-d) Choice fecundity assay results. Data are mean ± SD. * *P* < 0.05, χ2 test. Statistical analysis of chlorophyll content (e), leaf area (f and g) and leaf number (h) in different *A. thaliana* genotypes used in choice fecundity assays. Data are mean ± SD. * *P* < 0.05. One-way ANOVA with Tukey’s Multiple Comparison test for (e and h). Student’s *t*-test for (f and g).

## Notes

### Competing Interest Statement

The authors have declared no competing interest.

